# Actin conformation dynamics precede force generation in cytotoxic T lymphocytes

**DOI:** 10.1101/2025.05.15.654337

**Authors:** Adam M Rochussen, Aybaran O Kebabci, Alex K Winkel, Kristian Franze, Daniel A Fletcher, Gillian M Griffiths, Anna H Lippert

## Abstract

Cytotoxic T lymphocytes (CTLs) are cells of the adaptive immune system that are able to recognise and kill cancerous or virally infected target cells. The biochemical basis of CTL-mediated killing has been examined in great detail. Recent work has underscored the importance of physical forces in T cell function too, but how CTLs sense and exert force during migration and killing is not fully understood. Here, by expressing actin conformation probes based on the CH domain of utrophin, we directly visualize regions of altered F-actin conformation in primary CTLs during migration and killing. By combining these probes with traction force microscopy, we correlate external force with the internal cytoskeleton. We show that actin conformation is regulated upstream of force production, and that force exertion at the cytotoxic immune synapse temporally follows actin conformation dynamics. Our work offers novel insight into the relationship between cellular force production and F-actin conformation in important primary immune cells.

## Introduction

Cytotoxic T lymphocytes (CTLs) are important cells of the adaptive immune system responsible for the specific elimination of virally-infected and oncogenic cells presenting foreign peptides that can be recognised by the T cell receptor (TCR). These are highly mechanosensitive cells, encountering a wide range of stiffnesses from the soft thymus to stiffer inflamed cells ^1^. CTLs depend on force transduction in multiple different contexts: (i) migration, (ii) activation, and (iii) CTL-mediated killing ^2^.

T cells exhibit durotaxis, that is, enhanced migration on and towards stiffer substrates 3, likely mediated by actin-based adhesion modules, as has been shown in other cell types ^4^. Since inflammation increases the stiffness of antigen-presenting cells (APCs) 5, this durotaxis likely aids in the targeted migration of T cells during infection. Force-sensing and force-exertion go hand-in-hand for migratory cells, potentiating their migration through a range of tissues of differing stiffnesses ^6, 7^. For example, it is thought that the polymerisation of branched actin in leading-edge lamellipodia actively pushes cells forwards under certain modes of migration ^8^.

After migration, activation of T cells is also highly stiffness dependent. T cell activation, which occurs downstream of TCR ligation, is modulated by co-stimulatory receptors, cytokine signalling, and metabolic inputs ^9^. Recently, the physical stiffness of the APC and the transduction of forces have also emerged as essential factors, with stiffer APCs activating T cells better ^5, 10, 11^. Even high concentrations of high affinity ligands for TCR cannot induce T cell activation if the opposing surface is too soft ^12^. Notably, this property of T cell activation is one that is potentially exploited by cancerous cells, with softness correlating with metastatic potential and evasion of CTL-mediated killing^13–16^.

T cells not only sense force, but they also exert force, particularly at the immune synapse where forces are thought to play an important role in CTL-mediated killing ^17–19^. Notably, much mechanobiology of the immune synapse has depended on tractable immortalised T cell lines, such as Jurkat T cells, yet key differences between these model systems and primary T cells, as well as differences between CD4+ and CD8+ T cells, have been highlighted by others ^11, 20, 21^.

The actin cytoskeleton underpins the ability of T cells to generate forces, with myosin motors and varied actin polymerisation machineries all playing a role ^22, 23^. F-actin is highly dynamic at the cytotoxic immune synapse. Initial contact and T cell activation results in a characteristic ring of F-actin, with a depleted zone in the centre where TCR signalling, centrosome polarisation, and cytolytic protein secretion occurs ^24–26^. Retrograde actin flow and actomyosin arcs work to concentrate transmembrane receptors into the centre of the synapse, sustaining the signalling environment ^27^. Over time, the F-actin recovers across the synapse, blocking further secretion and marking the termination of the synapse before the cell detaches ^28^.

How the actin cytoskeleton exerts and responds to force is a subject of continued interest. One hypothesis posits that filament twist, altered by compression/tension, can affect the binding of actin-binding proteins, and thus transduce a physical force into a biochemical signal ^29^. Visualizing different F-actin conformations in living cells is possible via genetically encoded actin-binding proteins fused with fluorescent proteins, with different probes binding to the cytoskeleton in varied ways ^30^. For example, tandem calponin homology domains of the actin-binding protein utrophin (“utrn-CH”) is known to preferentially bind to F-actin near the uropod of migrating primary CD4^+^ T cells ^31, 32^, whilst the seventeen-residue polypeptide that constitutes LifeAct binds evenly across the length of migrating cells but fails to bind highly twisted cofilin-bound actin rods ^33, 34^. Structural analysis of utrn-CH revealed F-actin binding across three monomers, supporting its potential sensitivity to actin filament twist ^35^. Previously, variants of utrn-CH with altered F-actin-binding properties were developed, including the Q33A, T36A, K121A mutant (“utrnLAM”), which is enriched in lamellipodial networks in living cells ^36^. In a neutrophil-like cell line, simultaneous expression of the wildtype utrn-CH (“utrnWT”) and utrnLAM showed differential localisation along the front-rear axis during migration ^36^. These utrn probes could be easily displaced by physiologically relevant concentrations of actin binding proteins *in vitro*, implying they could be used as a ratiometric reporter of actin filament conformation within cells without impacting function, although this has not been tested ^36^.

Here, we optimize these probes for expression in primary CTLs and demonstrate that they exhibit biased localisation during migration without affecting migration speed or CTL function. We perform traction force microscopy (TFM) on CTLs expressing utrn-CH-based actin conformation probes during both migration and synapse formation. Inhibition of non-muscle myosin IIa, formins, or Arp2/3 is shown to disrupt force generation and actin structures, but not biased probe localisation, indicating upstream regulation of actin conformation in cells. Dynamic changes to actin conformation at the cytotoxic immune synapse, both on TFM gels and in CTL-target conjugates, revealed a compression-associated conformation dominating the early cytotoxic immune synapse, and a tension-associated conformation ensuing later on. Notably, force production over time mimicked the pattern of compression-associated F-actin conformation at the synapse, but with a time lag of ∼60 seconds. Thus, dynamical actin conformation is both regulated upstream of force production and temporally precedes force exertion in CTLs. Our work expands understanding of forces generated by T cells by focusing on primary CTLs (rather than cell lines) by presenting novel insight into actin conformation as it correlates with force production during both migration and killing.

## Results

### Actin conformation probes exhibit biased localisation in migrating primary CTLs and do not affect cell function

To generate equal expression of utrnWT and utrnLAM probes in primary CTLs, we combined separate constructs into a single plasmid, using a P2A sequence ^37^, and optimized fluorescent protein choice such that photobleaching would not significantly impact the fluorescence ratio between the two probes when imaging over time (supplementary fig. 1). Imaging CTLs transiently expressing these probes (“utrnP2A”) crawling on glass functionalized with the cell surface adhesion molecule ICAM-1 revealed a biased localisation, with a relative enrichment of utrnLAM versus utrnWT at filopodial tips and lamellipodia (fig. 1a, supplementary movie 1). This indicated varied actin filament conformation across the cytoskeleton of migrating CTLs and was consistent with what has been shown in immortalised neutrophil-like cells ^36^.

**Figure 1.**
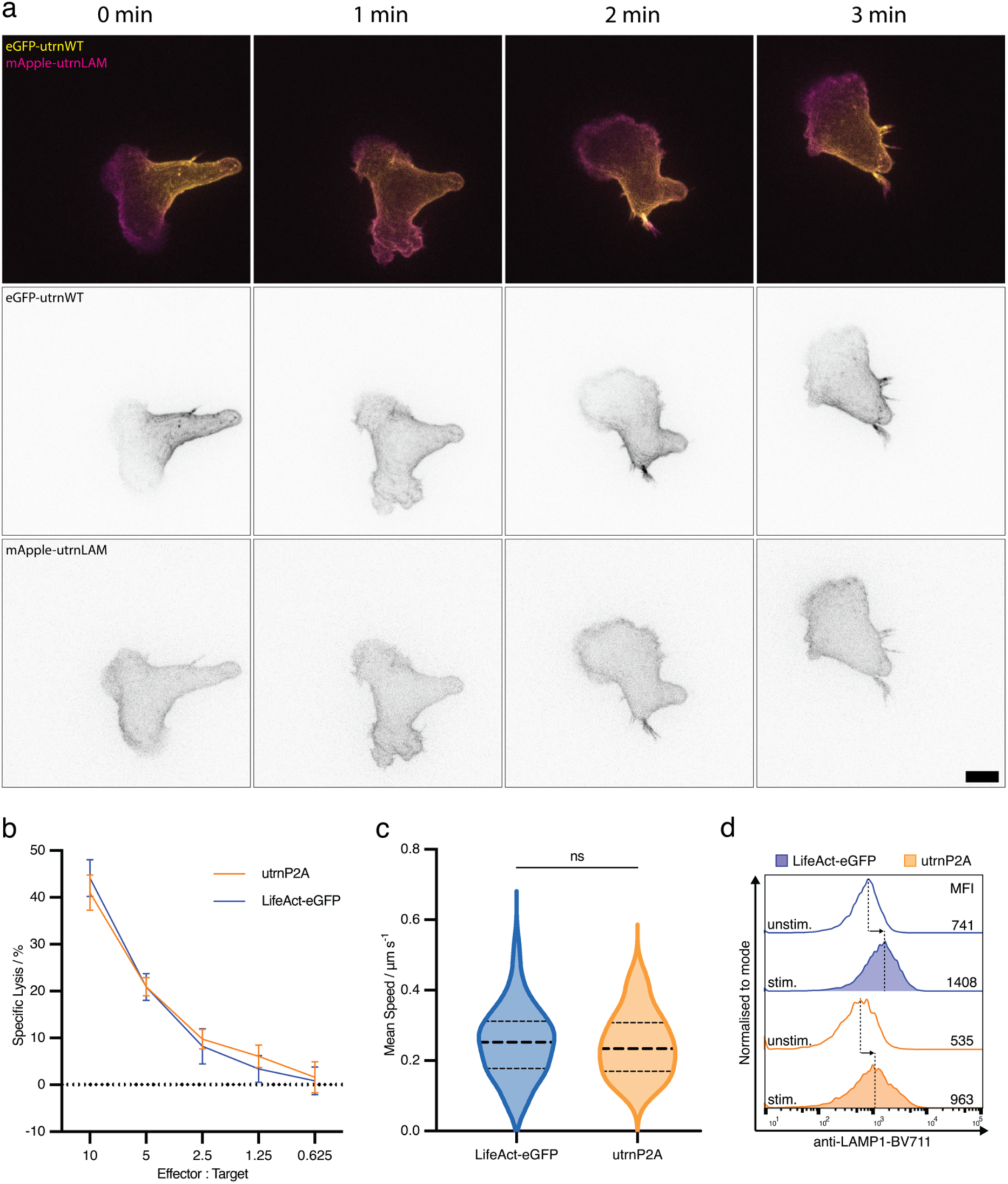
Actin conformation probes do not affect CTL function. a: Time-lapse spinning disc confocal microscopy of an OTI CTL crawling on ICAM1-coated glass. The cell is expressing eGFP-utrnWT (yellow in composite) and mApple-utrnLAM (magenta in composite) from the utrnP2A construct. Images are maximum intensity projections. Separate channels are shown in inverted greyscale as indicated. Scale bar represents 5 μm. Representative of at least 30 cells from at least five independent experiments. **b:** LDH killing assay comparing the ability of OTI CTLs expressing LifeAct-eGFP (blue) or utrnP2A (orange) to kill EL4 target cells at the indicated effector-to-target ratios. Representative of three independent experiments. **c:** Migration assay showing mean migration speed of OTI CTLs expressing LifeAct-eGFP or utrnP2A crawling on ICAM1-coated glass. Horizontal dotted lines represent median and interquartile range. Conditions were compared via unpaired t test (LifeAct mean = 0.2502 μm/s; utrnP2A mean = 0.2453 μm/s; p=0.7368). Representative of three independent experiments. **d:** Degranulation assay based on cell surface exposure of secretory lysosome protein LAMP1 in response to TCR activation. Median fluorescence intensity for anti-LAMP1-BV711 in each case is given. The rightwards shift upon stimulation is comparable between LifeAct-eGFP (blue) and utrnP2A (orange). Representative of three independent experiments.

We next sought to validate that CTL function was unaffected by their expression. Using LifeAct-eGFP as an alternative commonly-used F-actin probe for comparison ^33^, we found no detrimental impact of utrnP2A expression on CTL-mediated killing, degranulation, or migration speed (fig. 1b-d). Thus, utrnP2A could be reliably used to label areas of altered actin conformation in primary CTLs without affecting their function.

### Specific actin conformations are required but not sufficient for migratory force

To visualize force exertion by CTLs, we turned to traction force microscopy (TFM). Adapting a previously published protocol ^38^, we synthesised polyacrylamide hydrogels by polymerising an acrylamide mixture directly onto pre-cleaned and treated commercially available live imaging dishes with microbead imbedding via inverted polymerisation (supplementary fig. 2a, see Methods). Atomic force microscopy was used to measure gel stiffness and validate intra- and inter-batch consistency (supplementary fig. 2b-c). The resulting hydrogels exhibited an average Young’s modulus of 2.17 kPa, which matches the stiffness of a typical cancerous APC ^39^. Microbead distribution was even and appropriately spaced for high resolution TFM of primary CTLs (supplementary figure 2d).

Coating the TFM gels with ICAM-1, we seeded primary CTLs expressing utrnP2A and imaged their migration over time with a spinning disc confocal microscope. This revealed that biased localisation of actin conformation probes was maintained on soft hydrogels (supplementary movie 2, fig. 2). Using an established MATLAB script ^40^, we analysed the bead displacement and calculated traction force exerted by CTLs on the TFM gel (fig. 2). We found strong (∼ 6 Pa), forward-directed traction forces at the leading edge of the crawling CTLs, consistent with the notion of forceful branched F-actin polymerisation in lamellipodia during this type of migration ^8^.

**Figure 2.**
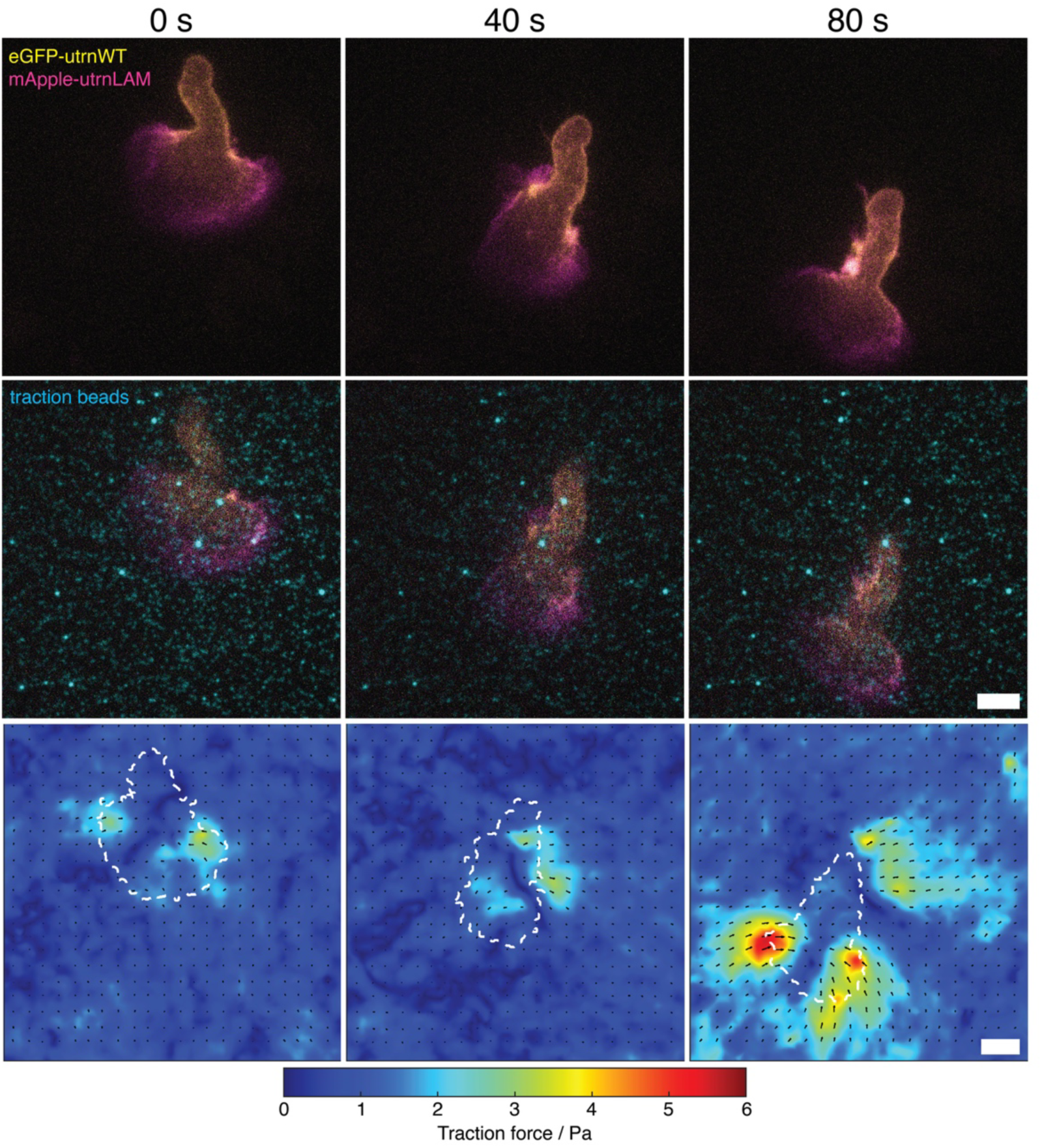
Biased localisation of actin conformations in CTLs migrating on soft hydrogels. Maximum intensity projection (top), single confocal slice in the plane of the TFM gel (middle), and traction force maps (bottom) of an OTI CTL, expressing utrn probes (eGFP-utrnWT, yellow; mApple-utrnLAM, magenta) crawling on an ICAM-coated TFM gel (AF647 microspheres, cyan) with 2.17 kPa Young’s modulus. Scale bars represent 5 μm. Traction force is indicated by colour scale between 0 (blue) and 6 Pa (red), with directional arrows shown. Representative of 40 cells across seven independent experiments.

To understand whether force generation is dependent on actin conformation or vice versa, we perturbed the ability of CTLs to produce forces via three distinct pathways: (i) paranitroblebbistatin (pNB) was used to inhibit non-muscle myosin IIa, a motor protein known to play a role in maintaining polarity of migrating cells, maturation of adhesion foci, and enhancing leading edge protrusion ^41–43^. (ii) CK666 was used to inhibit the Arp2/3 complex, which drives membrane protrusion during migration via actin branching, especially in lamellipodia. Arp2/3 is known to be mechanosensitive 44,45. (iii) SMIFH2 was used to inhibit formins, which are a group of proteins sharing formin homology domains that are responsible for linear actin polymerization ^46^. Formins are a diverse group of actin polymerisation proteins that have been linked to both migration and immune synapse formation ^27, 47–49^.

CTLs expressing utrnP2A were treated with each drug, seeded onto ICAM-1-coated TFM gels as before, and imaged over time. In DMSO-treated CTLs, lamellipodia and filopodia can be clearly seen, similarly to when migrating on glass (fig. 3a, supplementary fig. 3a, supplementary movie 2). With pNB treatment, leading edge lamellipodia are still present, but the rear of the cell is rounded and misshapen (fig. 3a, supplementary fig. 3a, supplementary movie 3), as has been observed by others with blebbistatin-treated cells ^50^. CK666-treated CTLs were unable to form lamellipodia, but instead formed filopodial protrusions at the leading edge as seen previously ^51^, with enhanced relative utrnLAM binding (fig. 3a, supplementary fig. 3a, supplementary movie 4). Finally, formin inhibition abrogated filopodia formation (fig. 3a, supplementary fig. 3a, supplementary movie 5). In each instance, regions of F-actin with biased probe localisation were persistently observed, despite an inability to maintain cell shape, form lamellipodia, or form filopodia respectively. Thus, biased localisation of F-actin conformation is regulated upstream or independently of actin-network structure.

**Figure 3.**
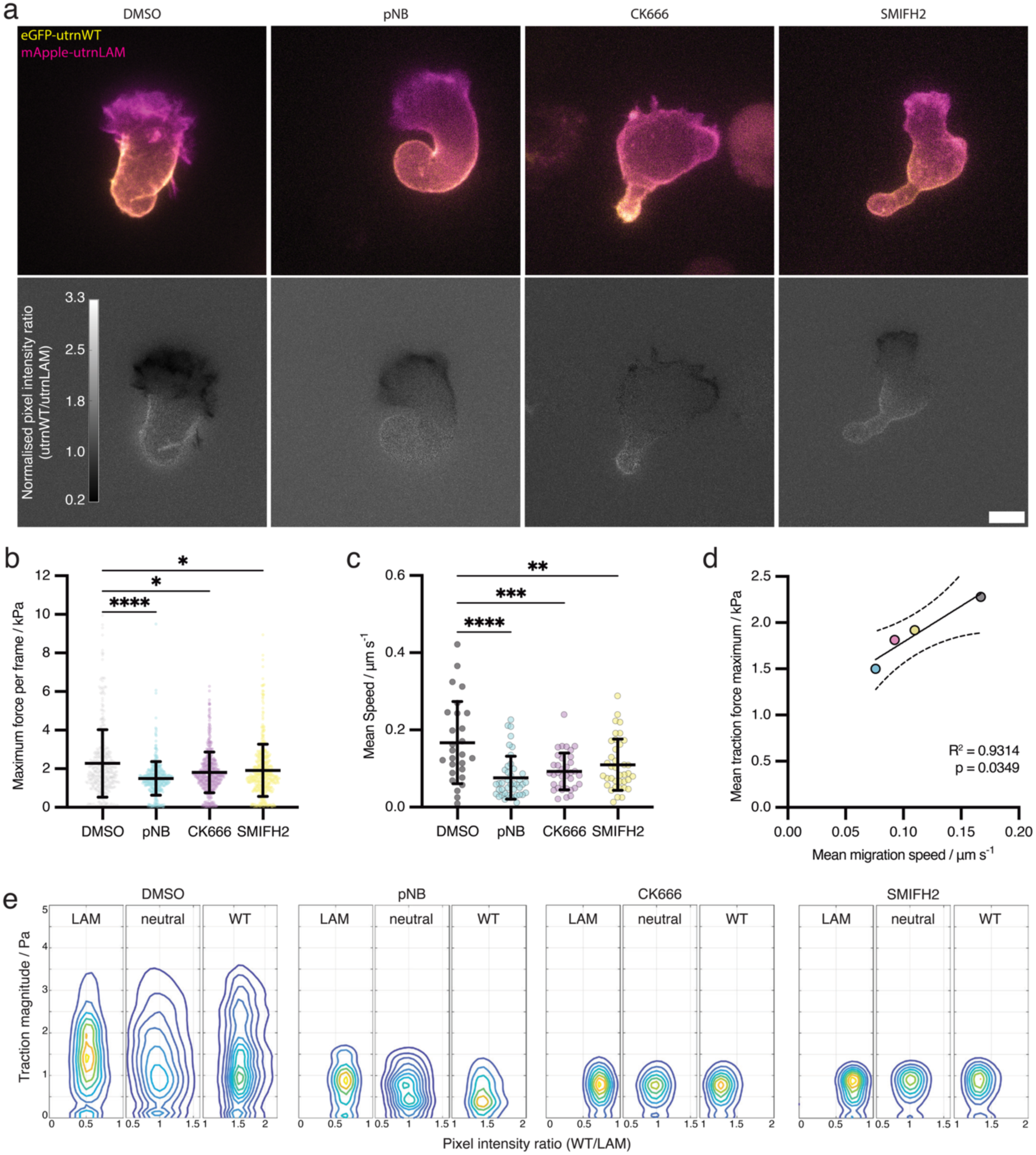
Perturbation of cytoskeletal machinery reduces migratory speed and force, but does not prevent biased probe localisation. **a**: Maximum intensity projections of single timepoints from timelapse spinning disc confocal microscopy of CTLs crawling on ICAM1-coated TFM gels. CTLs are expressing utrnP2A (eGFP-utrnWT, yellow, and mApple-utrnLAM, magenta). Ratiometric images are shown on the bottom row, where a low value (dark) indicates utrnLAM dominance. Separate channels are shown in supplementary fig. 3. CTLs were treated with the indicated drugs. Scale bar represents 5 μm. **b**: Comparison of maximum traction force per frame per cell between different drug treatments. Bars represent mean and standard deviations. Conditions were compared via Kruskal-Wallis ANOVA with Dunn’s multiple comparisons test: DMSO (mean = 2.28 Pa, n = 369) vs pNB (mean = 1.50 Pa, n = 420) p < 0.0001; DMSO vs CK666 (mean = 1.81 Pa, n = 492) p = 0.0142; DMSO vs SMIFH2 (mean = 1.92 Pa, n = 465) p = 0.0128. **c**: Comparison of migration speed on TFM gels. Mean cell speed was analysed using the TrackMate plugin in ImageJ and calculated in μm per second. Data points represent tracks from single cells. Mean and standard deviation are shown. Conditions were compared via one-way ANOVA with Dunnett’s multiple comparisons test: DMSO (mean = 0.1671 μm/s, n = 27) vs pNB (mean = 0.076 μm/s, n = 42) p < 0.0001; DMSO vs CK666 (mean = 0.0925 μm/s, n = 35) p = 0.0001; DMSO vs SMIFH2 (mean = 0.1099 μm/s, n = 38) p = 0.0036. Data representative of three independent biological replicates. **d**: Correlation between maximum traction force per frame and mean migration speed per track. Data points are the means from across all data points and all experiments for each of the four drug treatments (DMSO = grey, pNB = cyan, CK666 = magenta, SMIFH2 = yellow). Simple linear regression was performed with a best-fit line shown (𝑦 = 7.811𝑥 + 1.008) in solid line and 95% confidence intervals in dotted curves. R^2^ and p-values are shown. **e**: Pixel distribution of traction magnitude segregated by probe ratio from all timepoints of representative cells in each drug treatment condition. “LAM” shows traction force at pixels with a utrnWT/utrnLAM ratio smaller than one standard deviation below 1. “neutral” shows the same for ratios between one standard deviation below and one standard deviation above 1. “WT” shows the same for ratios greater than one standard deviation above one. All data representative of three independent experiments with 21 (DMSO), 14 (pNB), 17 (CK666), and 19 (SMIFH2) cells each across 12 frames of time lapse imaging.

Analysing the force produced by migrating CTLs treated with our suite of inhibitors revealed a significant loss of migratory forces with all drug treatments, with the most significant from pNB treatment (fig. 3b). Migration speed was similarly impacted by drug treatments, and force production and migration speed were found to be strongly positively correlated in migrating CTLs (fig. 3c, d).

As with untreated cells, forward-directed traction forces from the utrnLAM-dominant lamellipodia were observed with DMSO-treated cells (supplementary fig. 3d). To determine the relative contributions of the different actin conformations to force production, we created a ratiometric channel (fig. 3a) and segmented cells based on the ratio of probe expression in each pixel. Force production was found to predominantly occur in utrnLAM-dominant areas of the cytoskeleton of DMSO-treated cells, and, whilst regions of utrnLAM or utrnWT dominance persisted with drug treatments, this actin-conformation-correlated force production was abrogated with all drug treatments (fig. 3e, supplementary fig. 3b,c).

Taken together, these data show that, on ICAM1-coated surfaces, migratory forces and speed depend on non-muscle myosin IIa, Arp2/3, and formins, with none being dispensable. Further, inhibition of these cytoskeletal elements and their characteristic actin reduces force production specifically in utrnLAM-dominant areas of F-actin, without impacting the biased localisation of actin conformation. This suggests that altered actin conformation is generated by the cell upstream of specific actin network structures and force production.

### Force-associated actin conformation precedes force exertion at the immune synapse

Next, we turned to the immune synapse, where a high degree of force exertion has been observed ^52, 53^. We imaged primary CTLs expressing utrnP2A coming into contact and forming synapses with TFM gels functionalized with anti-CD3ε. A distinct spatiotemporal pattern of utrnWT and utrnLAM probe distribution was consistently observed. The initial contact included F-actin predominantly bound by utrnLAM that quickly depleted across the centre, forming an actin ring (t = 0 s, fig. 4a, supplementary fig. 3e, supplementary movie 6). As the synapse progressed, utrnWT-bound F-actin was observed accumulating inside the outer ring of utrnLAM-dominant F-actin (t = 100 s, t = 200 s), and this proceeded to recover across the centre of the synapse as the synapse matured (t = 300 s, fig. 4a, supplementary fig. 3e). Direct observation of bead displacement showed an inwardly directed force by the CTLs on the TFM gel (fig. 4b). This “pinching” by the actin-rich perimeter of the synapse is consistent with 2D TFM of Jurkat T cells ^52^.

**Figure 4.**
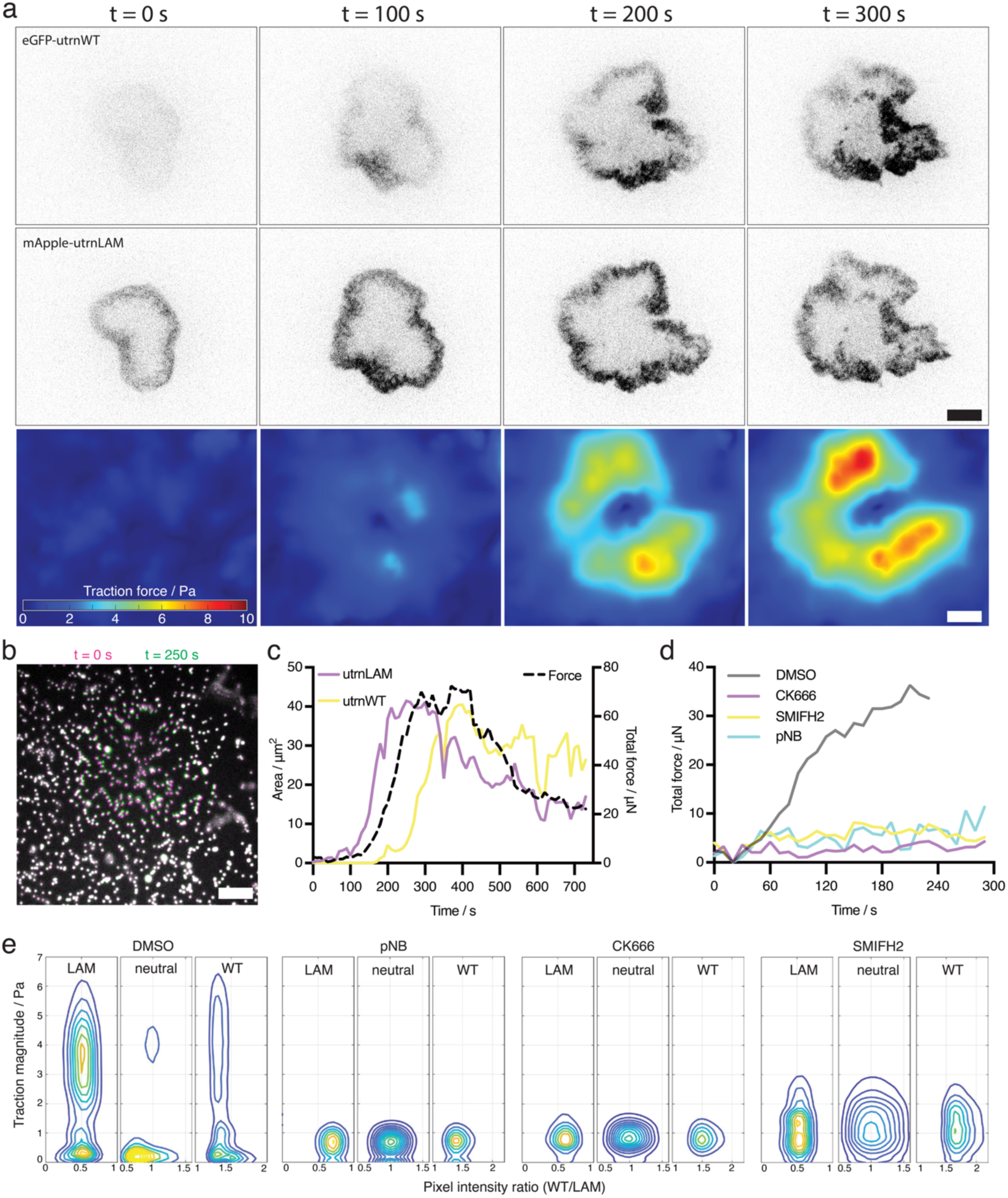
Dynamic changes to actin conformation at the cytotoxic immune synapse correlate with and precede force. **a**: Single confocal slices of a representative primary CTL, expressing utrn probes (merge images shown in supplementary fig. 3) forming a synapse with an anti-CD3ε-coated TFM gel. The first timepoint shown is the point of initial contact between CTL and gel. Traction force maps for each of the four time points depicted are also shown (bottom row), with force in Pa. **b**: Overlaid images of traction force beads from frame 1 (magenta) and frame 25 (green) relative to initial contact of the cell with the gel, relating to a. Where no bead movement has occurred, co-localisation results in white beads. Where there has been movement, the colours are seen separately. **c**: Analysis of probe area and traction force, relating to a. Left axis: area of eGFP-utrnWT (yellow) and mApple-utrnLAM (magenta) over time during artificial synapse formation. Areas were calculated by performing automatic Otsu thresholding on the time-stack per channel, and applying it to each frame. Right axis: total force production over time in μN. **d**: Total force production, in μN, over time by representative CTLs, expressing utrnP2A, treated with cytoskeletal inhibitors, forming synapses with TFM gels. Scale bars represent 5 μm. **e**: Pixel distribution of traction magnitude segregated by probe ratio for all timepoints from representative cells in each drug treatment condition (relating to d). All data are representative of 15 cells across three independent experiments.

Primary CTLs exerted significant traction force at the synapse, peaking at ∼ 10 Pa by 300 s (fig. 4a, supplementary movie 7). By plotting total force production and probe area over time, we determined that force correlated with utrnLAM distribution with a ∼50-100 s phase lag: The area of utrnLAM-bound F-actin rose steeply at 100 s, peaking near 40 μm^2^ at ∼250 s, before falling gradually as utrnWT-bound F-actin took over. Concurrently, total force rose sharply beginning at ∼150 s, peaking above 60 μN after 300 s, before falling gradually (fig. 4c). CTLs treated with cytoskeletal inhibitors, as before, were unable to produce force at the immune synapse (fig. 4d), whilst regions of biased probe localisation persisted as in our migration experiments (fig. 4e). The phase lag of force after utrnLAM-bound F-actin is further evidence that actin conformation is regulated upstream of force production, as shown by our TFM data of migrating CTLs treated with drugs (fig. 3). These data, showing force production at the synapse is preceded by utrnLAM-associated F-actin conformation dominance, imply that dynamical actin filament conformation is necessary but not sufficient for force exertion at the synapse.

To test whether these striking dynamic changes to actin conformation at the cytotoxic immune synapse were also observed at cell-cell synapses potentiated by physiological TCR:pMHC interactions, we mixed target cells, presenting cognate antigen and expressing membrane targeted Electra2, with primary CTLs expressing utrnP2A. Time lapse 3D confocal microscopy showed a highly consistent pattern, with a utrnLAM-dominant lamellipodial leading edge making initial contact and forming an actin ring (fig. 5a, supplementary movie 8, 9). F-actin at the synapse was dominated by utrnLAM for the first ∼200 s of the synapse, before actin conformation was altered, seen by an increased binding of utrnWT, culminating in complete recovery of cortical actin at the synapse by 800 s (fig. 5a, supplementary movie 8, 9). The temporal dynamics of actin conformation between physiological CTL:target conjugates and artificial TFM synapses were directly comparable, validating the physiological relevance of our data.

**Figure 5.**
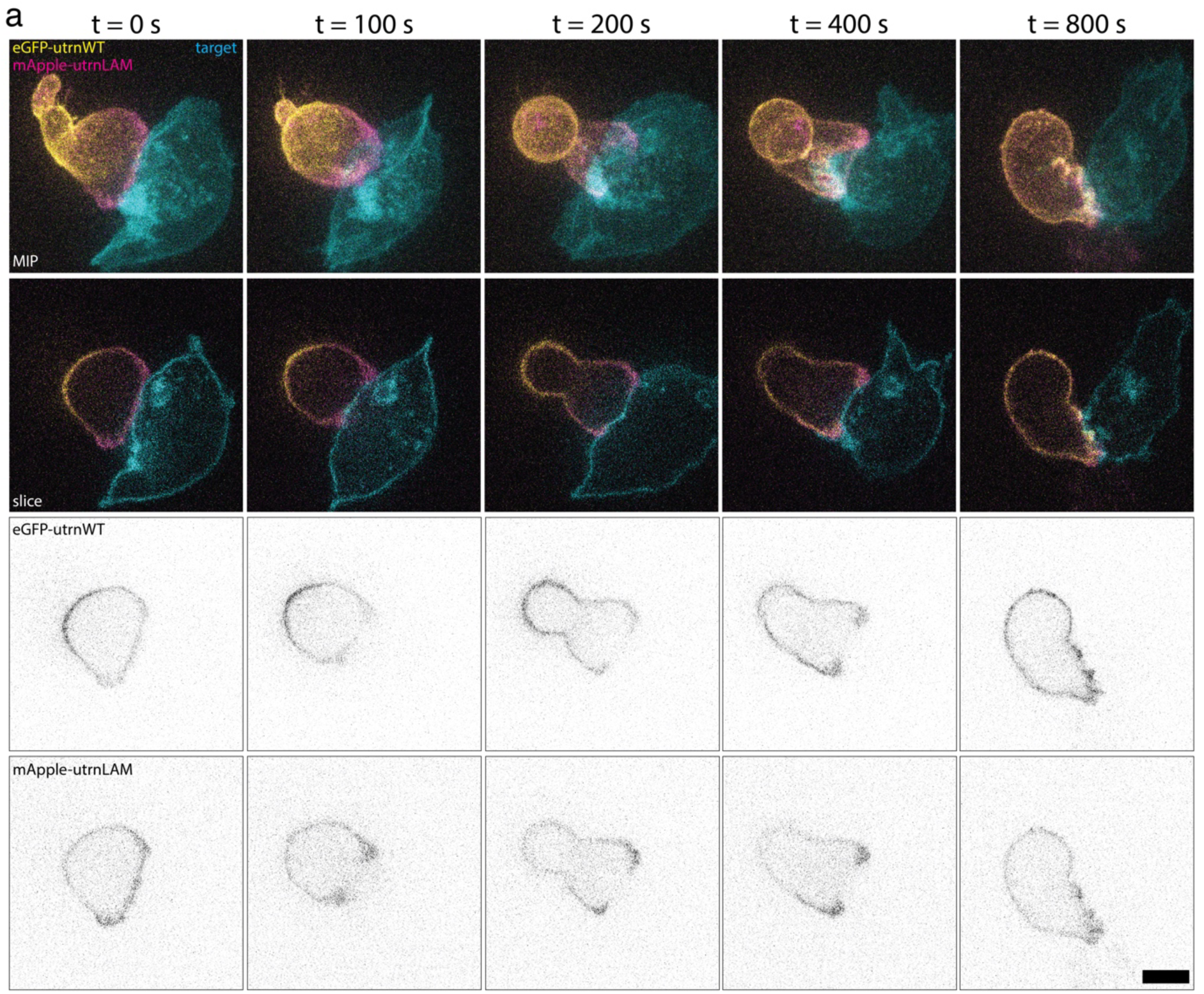
Dynamic changes to actin conformation at the physiological cytotoxic immune synapse. **a**: Timelapse spinning disc confocal microscopy of a representative primary OTI CTL, expressing utrnP2A (in merged images: eGFP-utrnWT in yellow, mApple-utrnLAM in magenta) forming an immune synapse with an EL4 target expressing membrane-targeted Electra2 (cyan) that had been pulsed with OVA257-264 peptide. The first timepoint shown is the point of initial contact between CTL and target. Maximum intensity projections are shown in the top row, and single confocal slices in the other three rows. Scale bar represents 5 μm. Representative of 15 cells from across three independent experiments.

## Discussion

By combining traction force microscopy of CTL migration and immune synapse formation with probes sensitive to actin conformation, we could observe the spatiotemporal correlation of actin conformations with force outputs by the cells. Force was found to be produced predominantly from regions of the cytoskeleton dominated by utrnLAM, whilst existence of these regions alone was not sufficient to produce force, as shown via treatment with cytoskeletal inhibitors. This force-associated F-actin (bound by utrnLAM) predominated the early immune synapse, being shortly followed by powerful force production. A shifting of the actin conformation to utrnWT-bound F-actin coincided with reduced force output and actin recovery across the centre of the immune synapse. Moreover, these spatiotemporal dynamics of F-actin conformation were consistent between traction force microscopy experiments and cell-cell cytotoxic immune synapses. Our work offers an exciting new parameter of the actin cytoskeleton that is important in both migration and killing by CTLs.

The binding of utrnLAM to leading-edge F-actin during migration correlated with a forward directed traction force, indicating compression of the actin filaments where utrnLAM binds. The idea that utrnLAM-bound F-actin is compressed, whilst utrnWT-bound actin is under tension, is consistent with the actin conformation changes we observed at the immune synapse, where an inward-directed pushing force was exerted by a utrnLAM-dominant F-actin ring, that then favoured utrnWT binding as cortical actin recovered across the centre of the synapse and force waned. Nonetheless, future work focusing on *in vitro* reconstitution will be required to elucidate the precise link between probe binding bias, actin conformation, and cytoskeletal forces in greater molecular detail.

Our use of cytoskeletal inhibitors reaffirmed prior work on the importance of non-muscle myosin IIa, Arp2/3, and formins in T cell migration and immune synapse formation. In addition, it allowed us to decouple actin conformation from force production. Our data here showed that differential probe localisation was maintained even while different actin-based structures (such as lamellipodia or filopodia) and force production were disrupted. This was surprising and implies that actin conformation is regulated upstream of force production, and is perhaps required for recruitment of actin binding proteins to manipulate F-actin in varied ways and exert force. Notably, since TFM can only measure forces external to cells, it is equally possible that exclusively internal forces (that do not transmit to the cell periphery) drive different filament conformations, in turn affecting utrn probe distribution.

The peak total force production we observed at the cytotoxic immune synapse (> 60 μN) is orders of magnitude higher than that reported for by Jurkat T cells on 2D TFM gels ^52^, potentially pointing to a difference between primary and immortalised cells. However, previous work by others involving 3D TFM of primary CTLs forming synapses with hydrogel beads also showed a much lower force output than we observed ^53^. In this instance, the TFM beads were much softer than our TFM gels (0.3 kPa vs 2.17 kPa), and indeed softer than most physiological target cells. The density of adhesive ligands (ICAM-1 or anti-CD3ε) may also contribute to this difference, since this is not standardised across studies. The lack of F-actin depletion across the centre of the immune synapse observed by Vorselen *et al.* is consistent with suboptimal T cell activation and may represent a failed cytotoxic immune synapse. Thus, the comparison of their data and ours suggests that high force production is essential for proper immune synapse function ^19^.

Since traction force microscopy is limited to *ex vivo* settings, fluorescence-based force probes that could be used *in vivo* are highly sought after to allow for biophysical studies to be performed in their proper physiological settings. Given the remarkable similarities between our 2D TFM and 3D cell-cell immune synapses, utrophin-based actin conformation probes have potential as a useful tool to study cellular biophysics in diverse *in vivo* systems, where TFM-based cellular biophysics is impossible.

## Methods

### Mice

C57BL/6 (B6)-OTI Rag2^−/−^ (B6.129S6-Rag2tm1Fwa Tg[TcraTcrb]1100Mjb) mice, referred to as OTI mice, were bred and housed in the University of Cambridge Facility. This research has been regulated under the Animals (Scientific Procedures) Act 1986 Amendment Regulations 2012 following ethical review by the University of Cambridge Animal Welfare and Ethical Review Body (AWERB). Spleens were obtained from male and female mice aged between 12 - 30 weeks.

### Plasmid design and cloning

For live fluorescence microscopy, the following plasmids were used:

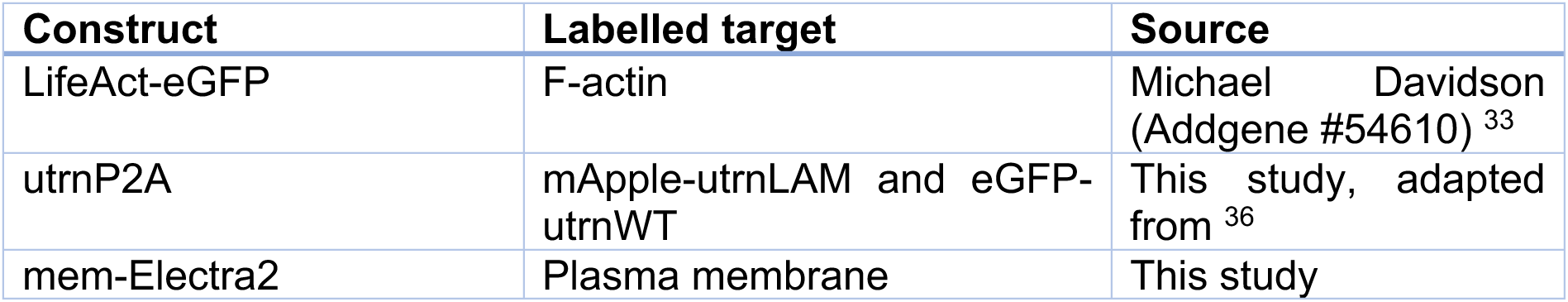

To design utrnP2A, the cDNA sequences of utrnLAM and utrnWT ^36^, were N-terminally fused with mApple and eGFP respectively. A GSG linker followed by a P2A sequence37 was inserted 3’ of eGFP-utrnWT (with stop codon removed), followed by mApple-utrnLAM. This construct was synthesised by Twist Biosciences (San Francisco, USA) with BamHI and NotI flanking sites, which were used to subclone into the pEGFP.N1 vector for transient expression in primary mouse T cells.

To design mem-Electra2, the first 20 amino acids of *Rattus rattus* neuromodulin ^54^ were fused to the N-terminus of blue fluorescent protein Electra2 via a short linker ^55^. The sequence was codon-optimised for *Mus musculus* and flanked with BamHI and NotI sites. The construct was synthesized by Twist Biosciences (San Francisco, USA) and subcloned into the pHR vector for lentiviral transduction.

### Cell culture

CTLs were from OTI mice, generated as previously described ^56^. Briefly, OTI splenocytes were stimulated with 10 nM OVA257-264 peptide (SIINFEKL) (Cambridge Bioscience) in mouse T cell medium: RPMI-1640 medium (SigmaAldrich, #1640) supplemented with 10% heat-inactivated FBS (LabTech, #FBS-SA), 50 mM β-mercaptoethanol (ThermoFisher, #31350010), 10 U/ml recombinant murine IL-2 (Peprotech, #212-12), 2 mM L-Glutamine (Sigma-Aldrich, #G7513), 1 mM sodium pyruvate (ThermoFisher Scientific, #11360070), and 50 U/ml penicillin and streptomycin (Sigma-Aldrich, #P0781). After three days, OVA257-264 was removed and cells were thereafter maintained in mouse T cell medium with daily replacement of medium via centrifugation and resuspension. Primary cells were incubated in a humidified atmosphere at 37 °C with 8% CO2.

Target cells for OTI CTL imaging experiments and killing assays were EL4 cells (ATCC: TIB-39; RRID: CVCL_0255) stably expressing mem-Electra2 (see below). Cell lines were maintained in DMEM (Sigma-Aldrich, #D5030) supplemented with 10% heat-inactivated FBS. Cell lines were incubated in a humidified atmosphere at 37 °C with 10% CO2.

### Lentiviral transduction

To generate EL4 target cells stably expressing mem-Electra2, Lenti-X 293 T cells (Takara, # 632180) were transfected with mem-Electra2 in the pHR backbone, pMD2.G (AddGene #12259), and pCMV ΔR8.2 (AddGene #12263) in a 5:1:4 ratio using TransIT reagent (Mirus Bio, #MIR 2704). Virus was harvested from the supernatant at 48- and 72-hours post-transfection, filtered, and concentrated using Lenti-X Concentrator (Takara, #631232). EL4 cells (ATCC: TIB-39; RRID: CVCL_0255) in logarithmic growth phase were resuspended in cell line medium at 4 x 10^6^/mL containing concentrated virus and polybrene at 1 μg/mL (Merck, #TR-1003-G). After 16 hours, the cells were diluted four-fold in cell line medium and cultured for three days before using FACS to isolate single-cell-derived clones based on Electra2 fluorescence.

### Nucleofection

OTI CTLs were nucleofected on day 4-6 post-stimulation from splenocytes. 5 x 10^6^ CTLs were washed in PBS via centrifugation (200 x g, 5 min) and resuspension, before final resuspension in nucleofection mix, comprised of 2.5 μg plasmid DNA and P3 Primary Cell Solution up to a total volume of 100 μL. Cells were electroporated in 100 µL Nucleocuvette™ vessels (Lonza, #V4XP-3024) with a 4D-Nucleofector® X unit (Lonza, #AAF-1003X) using pulse code DN-100. CTLs were immediately transferred, with micro-Pasteur pipette, to wells of a six-well plate containing 2 mL pre-warmed nucleofection recovery medium: calcium-free RPMI (US Biological, #R8999-02A), 5 % heat-inactivated FBS, 2 mM L-glutamine, 32 μM 1-thioglycerol (Merck, #M6145), 1.7 mM sodium pyruvate, 20 μM bathocuproine disulfate (Merck, #B1125). CTLs were left to recover at 37 °C with 8% CO2 for 6 hours, before topping up with 6 mL of pre-warmed mouse T cell medium.

### LDH killing assay

Killing of EL4 targets by OTI CTLs was determined by release of lactate dehydrogenase from lysed cells. Cells were seeded in round-bottomed 96-well plates at varied effector-to-target ratios, as indicated, in phenol red-free RPMI-1640 (ThermoFisher, #11835030) supplemented with 2 % heat-inactivated FBS, and 50 U/mL penicillin and streptomycin. Plates were centrifuged for 5 minutes at 300 x g prior to incubation at 37 °C with 8 % CO2 for three hours. Cell lysis was measured via CytoTox 96® Non-Radioactive Cytotoxicity Assay (Promega, #G1780). Briefly, after incubation the plate was centrifuged once more and supernatant transferred to a mirror-image flat-bottomed 96-well plate. Solubilised substrate mix (Promega, #G179A and #G180A) was added and incubated in the dark for 30 min, allowing for the enzyme-coupled reaction to proceed, resulting in conversion of tetrazolium salt into a red formazan product proportional to the number of lysed cells. After 30 min, absorbance at 490 nm was measured with a spectrophotometric plate reader. Specific target lysis was calculated by subtracting absorbance from medium-only control wells and comparing absorbance in wells containing effectors and targets with vs without OVA257-264 pulsing.

### Degranulation assay

CTLs were resuspended in fresh pre-warmed mouse T cell medium at 1 x 10^6^ cells/mL. Fluorescently conjugated anti-mouse CD107a (LAMP1) (ThermoFisher, clone 1D4B, #12-1071) was added to the medium at 2 μg/mL, before seeding CTLs onto flat bottomed plates that had been pre-coated with 1 μg/mL αCD3ε (‘stim.’) or PBS (‘unstim.’). Plates were incubated at 37 °C with 8% CO2 for the indicated time. Plates were then kept on ice and stained with a live-dead marker, before analysing with an Attune NxT flow cytometer (ThermoFisher). For LAMP1 cell surface exposure analysis, cells were gated on forward and side scatter to exclude debris and only include singlets, live-dead marker to exclude dead cells, and eGFP expression to exclude non-transfected cells.

### Live imaging assays

CTLs were nucleofected with fluorescent protein constructs 24 hours prior to imaging. Prior to imaging, cells were resuspended at 1 x 10^6^ cells/mL in imaging medium: phenol red-free RPMI supplemented with 10 % heat-inactivated FBS, 2 mM L-glutamine, 25 mM HEPES (ThermoFisher, #15630080), 50 U/mL penicillin-streptomycin.

For low-magnification migration speed assays: Imaging dishes (Mattek, #P35G-1.5-14-C) were pre-coated with 0.5 μg/mL ICAM-1 (R&D Systems, #796-IC) in PBS. 250 μL CTL suspension was added per dish and allowed 30 minutes at 37 °C with 8% CO2 for the cells to adhere and begin crawling. After this time, imaging medium was gently passed over the dish to remove non-adhered CTLs before imaging. Single confocal slices were captured with a 20x objective lens (Leica HC PL APO 20x) in the mid-plane of the cells every 5 seconds for 5 minutes per field of view. Only the FITC channel was acquired (LifeAct-eGFP or eGFP-utrnWT). Three fields of view were taken per condition per experiment.

For high-magnification live imaging, CTLs were resuspended in imaging medium and dropped onto functionalized glass or TFM hydrogel. In the case of artificial synapses, cells were immediately imaged to capture them coming into contact with the activating surface. In the case of imaging migrating CTLs, cell were left to settle, adhere, and begin to crawl for 30 min prior to imaging through a 100x objective lens (Leica HC PL APO 100x/1.40 OIL CS2). Confocal z-stacks with slices 0.8-1 μm apart were acquired over the whole volume of the conjugates every 10-30 seconds for up to 20 min.

For drug treatments, drugs were added to CTLs in imaging medium half an hour before imaging, and kept in throughout the assays. Working concentrations are given in the reagents table below.

In the case of imaging of conjugates, EL4 target cells expressing mem-Electra2 were pulsed with 1 μM OVA257-264 peptide for half an hour and then washed thrice by centrifugation and resuspension in cell line medium. Target cells were ultimately resuspended in RPMI (serum-free) at 1 x 10^6^ cells per mL, and 250 μL seeded onto imaging dishes that had been pre-coated with ICAM as before. After 10 minutes at 37°C with 8% CO2, imaging medium was gently passed over the imaging dish to remove non-adhered target cells. Nucleofected CTLs were dropped onto target cells and allowed 10 minutes to settle before imaging.

All live microscopy used an Andor spinning-disk confocal system (Revolution; Andor) fitted with a CSU-X1 spinning-disk unit (Yokogawa) via a DMi8 microscope (Leica). Samples were excited with a combination of lasers of 405, 488, 561, and 637 nm wavelength. Images were captured using an iXon Ultra 888 camera and Fusion software (Andor). Imaging was performed with cells at 37 °C, 5 % CO2 in an airflow sample chamber on the stage (OkoLab).

### Image analysis

Migration assay data was analysed with ImageJ software (NIH) using the TrackMate plugin. Cells were masked using a Laplacian of Gaussian detector, with an estimated diameter of 11 μm and threshold of 2.0. Migration was tracked between frames via simple LAP tracker, with 5 μm maximum linking distance and zero tolerance for gaps. Tracks with fewer than 10 spots were excluded from analysis. Mean track speed was exported and plotted in Prism software (GraphPad).

Traction force microscopy assays were analysed via MATLAB, customising a published script (GitHub: https://github.com/DanuserLab/u-inferforce) ^40^. For the TFM analysis of the bead movement, the slice of the confocal z-stack, where the surface of the gel was in focus, was selected. Frames were drift-corrected relative to each other by efficient subpixel registration, and bead displacement was calculated frame-to-frame. Using the empirically measured gel stiffness of 2.17 kPa, traction force was then calculated using Fourier Transform Traction Cytometry (FTTC), with the L-curve method employed for regularization parameter selection, using L-optimal criteria. For maximal force value per frame, pixel resolution was possible. To determine the probe ratio in the maximal force zone, a 30 px radius circle centred on the maximal force pixel was used to ascertain the average probe ratio in this area.

For TFM analysis based on utrn probe distribution, each frame was normalised to the maximum intensity pixel, before histogram-matching in each probe channel. For ratiometric cell segmentation, utrnWT/utrnLAM ratio was calculated and categorised as “utrnLAM dominant”, “utrnWT dominant”, or “neutral” based on a ratio value two standard deviations below 1, two standard deviations above 1, or between the two thresholds. For migrating cells, maximum intensity projections were used to create the ratiometric image. For synapses, only the bottom z-slice (in focus with the synapse) was analysed.

For analysis of utrn probe area over time at the synapse, we generated time-stacks of each probe in the plane of the gel and performed automatic Otsu thresholding (such that all timeframes were incorporated into the thresholding algorithm). Total area per probe for each frame was then calculated based on these thresholds.

Data from imaging of live conjugates were analyzed and processed for export in Imaris software (BitPlane). Image channels were pseudo-coloured for optimal visual contrast.

### Traction force microscopy gel preparation

Gel preparation was adapted from published work ^38^ as follows:

For glass preparation: Imaging dishes (Mattek, #P35G-1.5-14-C) were cleaned via three 5 min incubations with 100 % ethanol. After final aspiration, glass was left to air-dry completely. The glass was then silanized via 30 min incubation at room temperature with 0.5 % (3-aminopropyl)trimethoxysilane (APTMS) (Sigma-Aldrich, #281778) in ultrapure sterile water. APTMS solution was then aspirated and the glass rinsed with water thoroughly. The surface was prepared for gel adhesion by incubation for 30 min at room temperature with 0.25 % glutaraldehyde (Sigma-Aldrich, #354400). The glass was rinsed thoroughly with water again and left to airdry. Top coverglasses were prepared by cleaning 18 mm circular glass coverslips (VWR, #630-2200) via water bath sonication in 100 % ethanol for 10 min, then leaving to airdry on clean Kimwipe.

For gel preparation: Gel premix was created by adding 500 μL of 40 % acrylamide solution (SLS, #A4058) and 65 μL of N-hydroxyethyl acrylamide (Sigma-Aldrich, #697931) to an eppendorf tube and mixing. 65 μL was then removed (leaving 500 μL), and 250 μL of 2 % N,N’-methylenebisacrylamide solution (Sigma-Aldrich, #M1533) was then added and mixed by vortexing. To create TFM gels with an average stiffness of 2.17 kPa, 65 μL of gel premix was added to 425 μL of DPBS (Gibco, #14190144) and mixed. To this, 10 μL of dark red 0.2 μm FluoSpheres™ Carboxylate-Modified Microspheres (ThermoFisher, #F8807) was added. This mixture was vortexed and then sonicated in a water bath sonicator for 10 min, and subsequently de-gassed via vacuum desiccator for 4 min.

For TFM gel synthesis: To polymerize the gel, 1.5 μL of N,N,N’,N’- tetramethylethylenediamine (TEMED) (Sigma-Aldrich, #T9281) and 5 μL of 10 % (w/v) ammonium persulfate solution (Sigma-Aldrich, #A3678) were added to 500 μL of TFM gel mix and mixed by gentle pipetting. As quickly as possible, 3 μL was added to the centre of the prepared glass imaging dish. A prepared top coverglass was then inverted on top, causing the gel to spread to the edges of the 18 mm glass coverslip. The entire imaging dish was then inverted and left for 15 min at room temperature for the gel to polymerize. Flipping the dish back again, DPBS was added to cover the glass and left for 30 min. Then the top coverglass was removed gently with tweezers. Gels were treated with poly-D-lysine (Gibco, #A3890401) overnight at 4 degrees C. The gel could then be functionalized with a biologically relevant molecule.

For migration on TFM gels, poly-D-lysine was rinsed three times with PBS, then replaced with 50 μg/mL ICAM-1(R&D Systems, #796-IC) in PBS and incubated at 37 degrees C for 1 hour, before three PBS washed and cell mounting. For artificial immune synapses on TFM gels, the same process was performed but substituting ICAM-1 for anti-CD3ε (ThermoFisher, #14-0033-82) at the same concentration of 50 μg/mL. High concentrations were used to compensate for poor functional molecule adhesion compared to regular glass coverslips, ensuring saturation of the surface.

### Atomic force microscopy

AFM measurements were performed on a JPK CellHesion200 (Bruker), with a tipless cantilever (ArrowTL, NanoWorld) to which a 37 μm diameter polystyrene bead (PS-R-37.0 - microParticles Gmbh) had been glued. The spring constant of the cantilever, determined by thermal tuning, was 0.092 N/m, the setpoint 10 nN and the speed of the Z scanner was 10 μm/s. Force distance curves were analysed in the JPK Data Processing software (Bruker) where the Hertz-Sneddon Model was applied to calculate the Young’s Modulus, assuming Poisson’s ratio to be 0.5.

### Data presentation

Data was processed, statistical tests applied, and data plotted in Excel (Microsoft) and/or Prism (GraphPad). Flow cytometry data was plotted in FlowJo (BD). Schematic diagrams were produced in BioRender. Figures were collated in Illustrator (Adobe). Manuscript was processed in Word (Microsoft).

### Table of reagents

| DRUGS | Name | Target | Effect | Working concentration | Source | Catalogue number |
| --- | --- | --- | --- | --- | --- | --- |
|  | CK666 | Arp2/3 | Inhibition | 100 µM | SLS | 182515 |
|  | (S)-4'-nitro-blebbistatin | Non-muscle myosin II | Inhibition | 20 µM | Cayman Chemical | CAY24171 |
|  | SMIFH2 | Formin FH2 domain | Inhibition | 20 µM | Merck | 344092 |
| DYES | Name | Target | Use | Working concentration | Source | Catalogue number |
|  | LIVE/DEAD™ Fixable Yellow Dead Cell Stain | Dead cells | Flow | 1:400 (flow) | ThermoFisher | L34959 |
| ANTIBODIES | Target | Host species | Clone | Working concentration | Source | Catalogue number |
|  | CD3ε (mouse), activating | Syrian hamster | eBio500A2 | 1 µg/mL | ThermoFisher | 14-0033-82 |
|  | LAMP1 (mouse), BV711 conjugated | Rat | 1D4B | 1:200 (flow) | BioLegend | 121631 |

## Authorship

AHL and DAF conceived the study. AHL, AMR, GMG, and KF sourced funding for the study. AMR and AHL planned experiments. AMR and AHL performed TFM experiments. AKW performed atomic force microscopy measurements. AMR performed all other experiments. AOK performed all TFM analysis. AMR performed all other image analysis. AMR wrote the original manuscript. All authors reviewed and revised the manuscript.

## Acknowledgements

We thank Huw Colin-York for critical reading of the manuscript, and Andrew Harris and other Fletcher lab members for providing the original utrn probe plasmids. We thank Mark Bowen and Matthew Gratian at the CIMR microscopy core facility, and Reiner Schulte and Gabriela Grondys-Kotarba at the CIMR flow cytometry core facility for training, access to equipment, and assistance. We thank Yukako Asano for assistance with microscopy. This work was supported by Wellcome Trust grants [102163/B/13/Z], [217100/Z/19/Z], and [215899/Z/19/Z].

---

**Supplementary figure 1.**
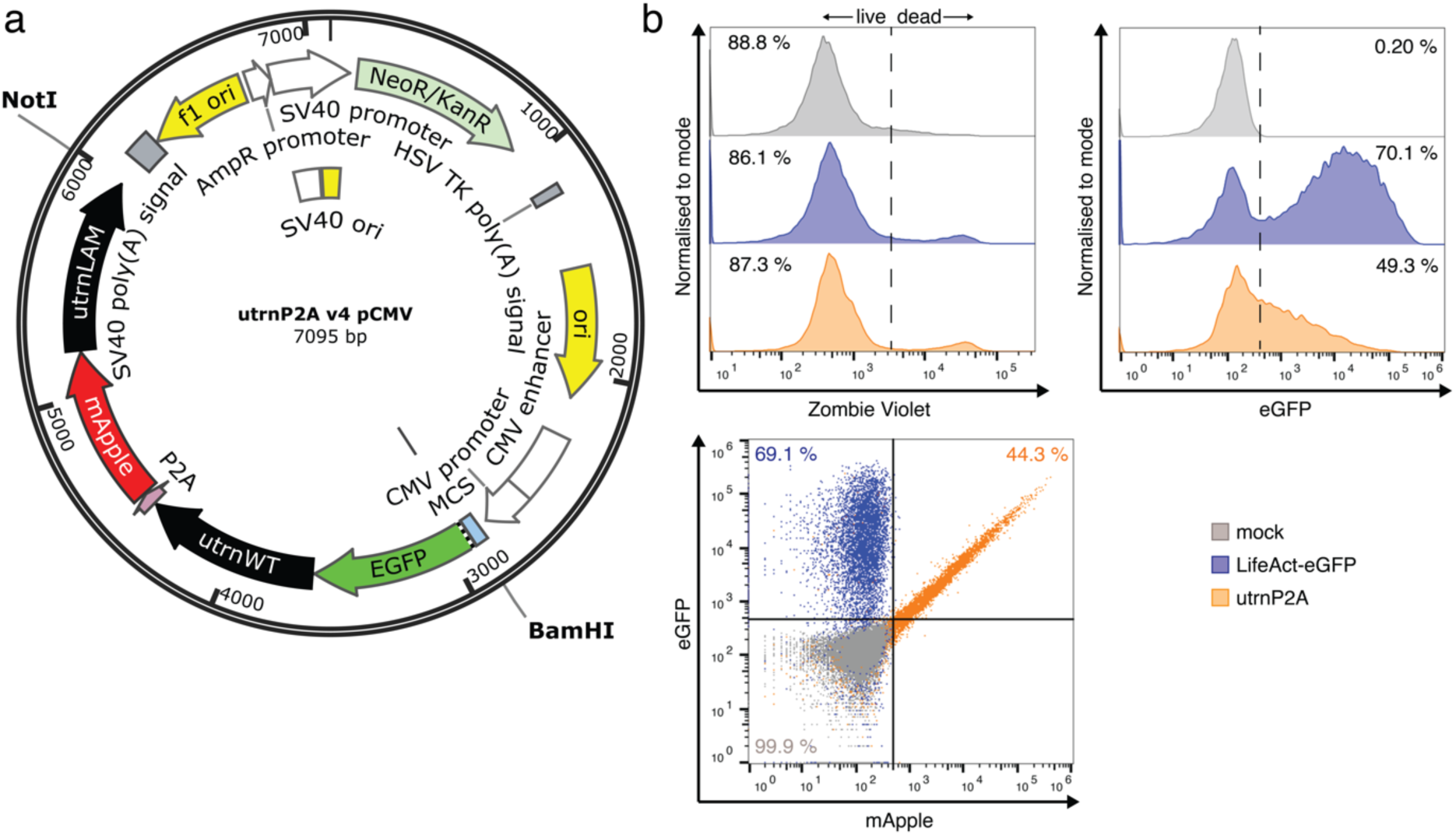
Expressing utrn probes in primary CTLs. a: Plasmid map for utrnP2A construct. Key features are labelled, including restriction sites used for subcloning (BamHI and NotI). **b:** Flow cytometric analysis of CTLs 24 hours after nucleofection with nothing (mock, grey), LifeAct-eGFP (blue), or utrnP2A construct (orange). Histograms depicting viability (top left), eGFP expression (top right), and dot plot comparing eGFP and mApple expression (bottom left). Representative of three independent experiments.

**Supplementary figure 2.**
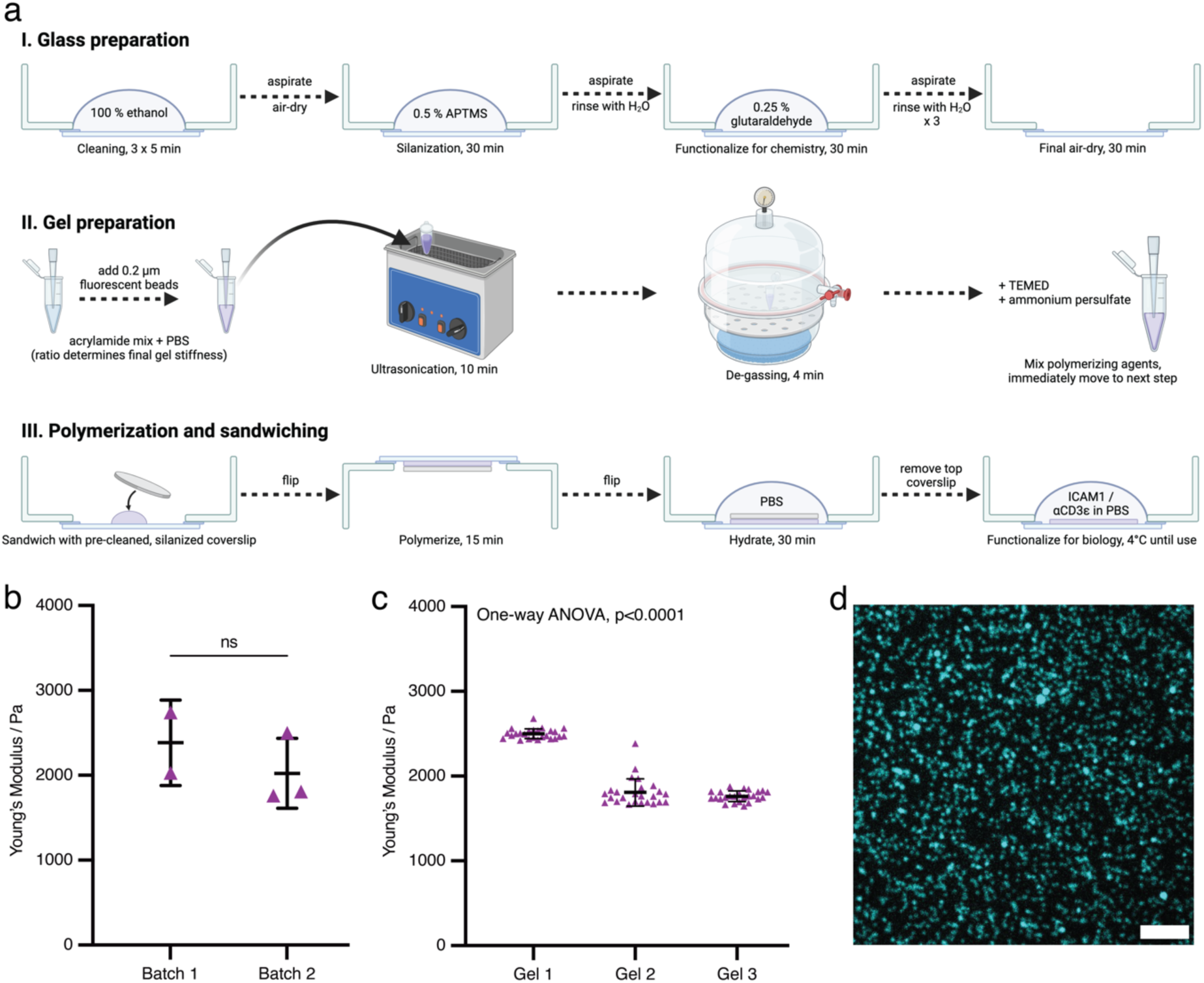
Synthesis and validation of traction force microscopy gels. a: Schematic diagram summarising the protocol for synthesis of traction force microscopy gels. Made in BioRender. **b-c**: Atomic force microscopy measurements of gel stiffness. Representative gels were selected from successive batches for AFM measurements. Twenty-five measurements were made per gel, sampling from the full area of the gel. **b:** Inter-batch consistency. Data points are means of 25 measurements per gel. Bars depict mean and standard deviation. There was no significant difference between batches (two-tailed t test, p=0.44). **c:** Homogeneity within gels. Measurements across each gel gave tightly clustered stiffness readings (ordinary one-way ANOVA, p<0.0001). **d:** Representative field of view of a TFM gel, imaged with a spinning disc confocal microscopy. Even distribution of dark red FluoSpheres™ (cyan) embedded beneath the surface can be seen. Scale bar represents 5 μm}

**Supplementary figure 3.**
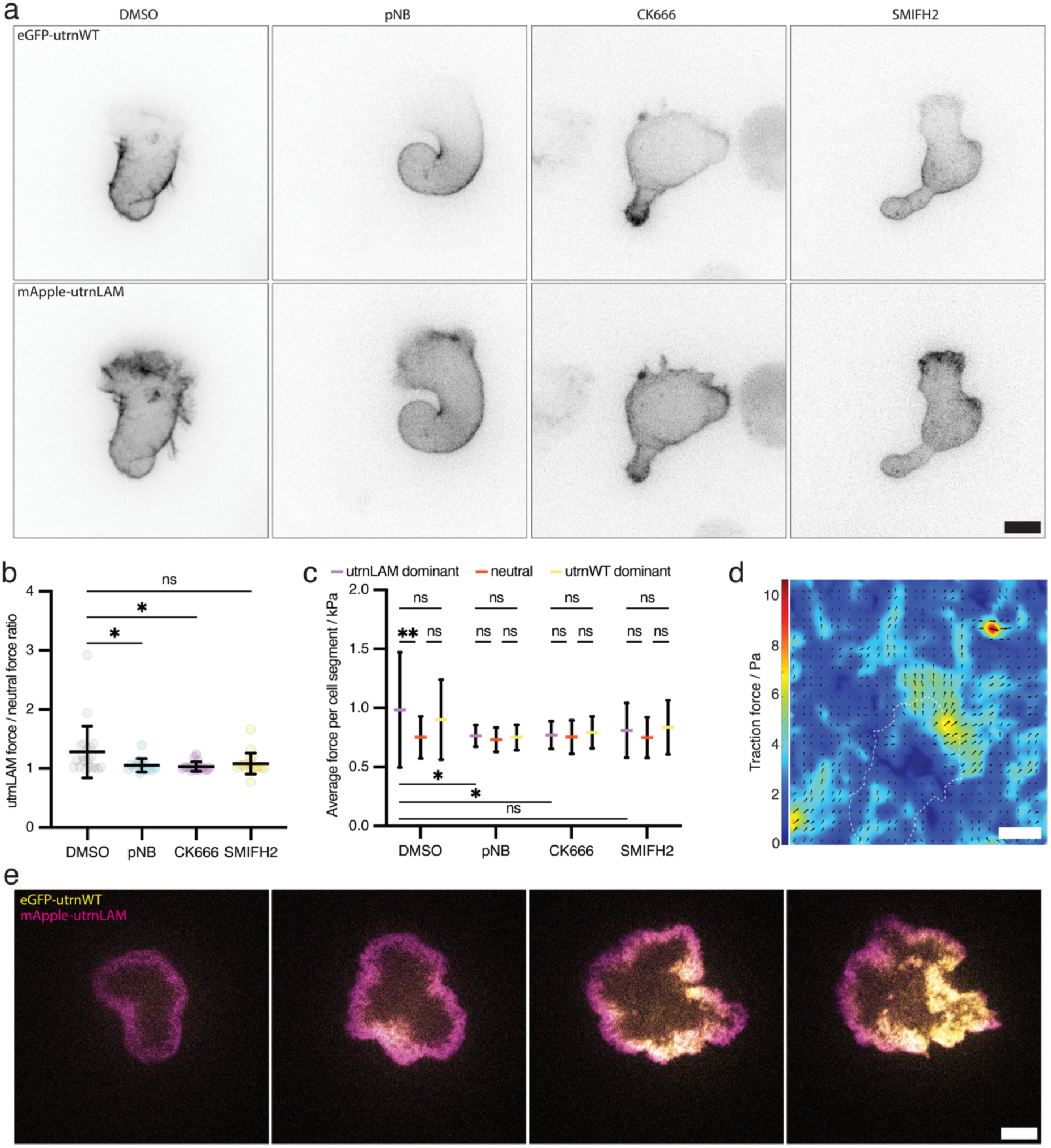
Abrogation of force production and actin network structures does not affect biased probe localisation. a: OTI CTLs expressing utrnP2A crawling on ICAM-coated TFM gels treated with the indicated drugs. Separated channel images in inverted greyscale relating to figure 3a. **b**: Ratio of average traction force value per movie between utrnLAM dominant and neutral cell areas between different drug treatments. Data points represent time lapse imaging series of individual cells. Bars represent mean and standard deviations. Conditions were compared via one-way ANOVA with Dunnett’s multiple comparisons test: DMSO (mean = 1.280) vs pNB (mean = 1.053) p = 0.0409; DMSO vs CK666 (mean = 1.031) p = 0.0146; DMSO vs SMIFH2 (mean = 1.081) p = 0.0549. **c**: Comparison of average traction force per movie per segmented cell area between different drug treatments. Cells were segmented into three areas based on ratio of utrn probe intensity. Average force production in each area across time lapse imaging was calculated. Bars represent mean and standard deviations. Conditions and cell areas were compared via two-way ANOVA and Šídák’s multiple comparisons test: utrnLAM dominant vs neutral DMSO, p = 0.0052; utrnLAM dominant DMSO vs pNB, p = 0.0466; utrnLAM dominant DMSO vs CK666, p = 0.0374; utrnLAM dominant DMSO vs SMIFH2, p = 0.1239. **d**: Representative traction force field with directional arrows for a DMSO-treated OTI CTL crawling on ICAM-1 coated TFM gel relating to Figure 3a. Scale bar represents 5 μm. Traction force is colour coded in Pa. Cell outline is shown in dotted white. **e**: An OTI CTL expressing utrnP2A (eGFP-utrnWT in yellow, mApple-utrnLAM in magenta) forming an immune synapse with an anti-CD3ε-coated TFM gel. Merged channel image relating to Figure 4a.

**Supplementary movie 1 - Actin conformation probes exhibit biased localisation in migrating CTLs.** Time-lapse spinning disc confocal microscopy of an OTI CTL crawling on ICAM1-coated glass. The cell is expressing eGFP-utrnWT (yellow) and mApple-utrnLAM (magenta) from the utrnP2A construct. Maximum intensity projections shown. Scale bar represents 5 μm. Time in mm:ss format. Representative of at least 30 cells from at least five independent experiments.

**Supplementary movie 2-5** - **Traction force microscopy of migrating CTLs expressing actin conformation probes.** Time-lapse spinning disc confocal microscopy of an OTI CTL, expressing utrn probes (utrnWT, yellow; utrnLAM, magenta) crawling on an ICAM-coated TFM gel (AF647 microsheres, cyan) (*E* = 2.17 kPa). Representative maximum intensity projections shown. Scale bar represents 5 μm. Time is given in seconds. Cells were treated with DMSO (Movie 2) (N = 3, n = 21), paranitroblebbistatin (Movie 3) (N = 3, n = 14), CK666 (Movie 4) (N = 3, n = 17), or SMIFH2 (Movie 5) (N = 3, n = 19).

**Supplementary movie 6 – Dynamic changes to actin conformation at the cytotoxic immune synapse.** Time-lapse spinning disc confocal microscopy of an OTI CTL, expressing utrn probes (utrnWT, yellow; utrnLAM, magenta) forming a synapse with an anti-CD3ε-coated TFM gel (AF647 microsheres, cyan) (*E* = 2.17 kPa). Cell channels (left) and drift-corrected bead channel (right) are separated to better visualize bead movement. Single confocal slice in plane with the gel surface was acquired. Scale bar represents 5 μm. Time given in mm:ss format. N = 3, n = 5.

**Supplementary movie 7 – Traction force at the cytotoxic immune synapse.** Traction force reconstruction relating to supplementary movie 6. Each frame is 10 seconds apart, and the image is 35 μm wide. Colour-coded traction force is in Pascals.

**Supplementary movie 8-9 – Dynamic changes to actin conformation at the physiological cytotoxic immune synapse.** Timelapse spinning disc confocal microscopy of a representative primary OTI CTL, expressing utrnP2A (eGFP-utrnWT in yellow, mApple-utrnLAM in magenta) forming an immune synapse with an EL4 target expressing membrane-targeted Electra2 (cyan) that had been pulsed with OVA257-264 peptide. Maximum intensity projections (Movie 8) and a single confocal slice through the centre of the synapse (Movie 9) are shown. Scale bar represents 5 μm. Representative of 15 cells from across three independent experiments.

